# StoHi-C: Using t-Distributed Stochastic Neighbor Embedding (t-SNE) to predict 3D genome structure from Hi-C Data

**DOI:** 10.1101/2020.01.28.923615

**Authors:** Kimberly MacKay, Anthony Kusalik

## Abstract

In order to comprehensively understand the structure-function relationship of the genome, 3D genome structures must first be predicted from biological data (like Hi-C) using computational tools. Many of these existing tools rely partially or completely on multi-dimensional scaling (MDS) to embed predicted structures in 3D space. MDS is known to have inherent problems when applied to high-dimensional datasets like Hi-C. Alternatively, t-Distributed Stochastic Neighbor Embedding (t-SNE) is able to overcome these problems but has not been applied to predict 3D genome structures. In this manuscript, we present a new workflow called StoHi-C (pronounced "stoic") that uses t-SNE to predict 3D genome structure from Hi-C data. StoHi-C was used to predict 3D genome structures for multiple, independent existing fission yeast Hi-C datasets. Overall, StoHi-C was able to generate 3D genome structures that more clearly exhibit the established principles of fission yeast 3D genomic organization.

## INTRODUCTION

Understanding the structure-function relationship of various biomolecules has been the foundation of molecular biology research for many years. Recently, the development of Hi-C (and related methods) has resulted in the generation of unprecedented sequence-level investigations into the structure-function relationship of the genome (Lieberman-Aiden *et al.* 2009; Belton *et al.* 2012; Belaghzala *et al.* 2017). Hi-C is able to detect regions of the genome that are "interacting" (*i.e.* in close 3D spatial proximity). Typically, this is done by mapping Hi-C sequence reads to a reference genome (Lajoie *et al.* 2015; Wingett *et al.* 2015; MacKay *et al.* 2018; MacKay and Kusalik 2019). This results in the generation of a whole-genome contact map which is a *N × N* matrix where *N* is the number of “bins” which represent linear regions of genomic DNA. Each cell within a whole-genome contact map records a count of how many times two genomic regions were found in close proximity within a population of cells (Lajoie *et al.* 2015; MacKay *et al.* 2018; MacKay and Kusalik 2019). This is more commonly referred to as an interaction count. Interaction counts are often normalized using methods like iterative correction and eigenvector (ICE) decomposition (Imakaev *et al.* 2012; Varoquaux and Servant 2019) to reduce inherent biases within Hi-C datasets (Yang and Jiang 2014; Li *et al.* 2015; Servant *et al.* 2015; Stansfield *et al.* 2018; Lyu *et al.* 2019; Spill *et al.* 2019). This normalization process results in fractional interaction counts also known as interaction frequencies.

Normalized whole-genome contact maps can be used to infer 3D genomic structure(s). This is known as the 3D genome reconstruction problem (3D-GRP) (Segal and Bengtsson 2015; MacKay and Kusalik 2019) or the 3D chromatin structure modelling problem (Zhang *et al.* 2013). For the purpose of this manuscript, we will be using the term 3D-GRP. A formal representation of the 3D-GRP is provided by MacKay and Kusalik (2019). Briefly, normalized interaction frequencies are converted into a set of pairwise distances (based on the inverse of the interaction frequency). This calculation uses the assumption that a pair of genomic regions with a small interaction frequency will be further away in 3D space than a pair of genomic regions with a higher interaction frequency (*Duan et al.* 2010; Fraser *et al.* 2010; Rousseau *et al.* 2011; Baù and Marti-Renom 2011, 2012; Hu *et al.* 2013; Ay *et al.* 2014; Lesne *et al.* 2014; Varoquaux *et al.* 2014; Sekelja *et al.* 2016; MacKay and Kusalik 2019). Each genomic bin’s (*x*, *y*, *z*) coordinates are then calculated using various optimization techniques (MacKay and Kusalik 2019).

Many of the existing tools for solving the 3D-GRP rely on multidimensional scaling (MDS) either completely or partially to predict and embed genomic structures in 3D space. MDS is known to have inherent problems when calculating embeddings from population-based, sparse, high-dimensional datasets (which are characteristics of Hi-C datasets) (Adhikari *et al.* 2016; Rieber and Mahony 2017). Alternatively, t-Stochastic Neighbourhood Embedding (t-SNE) has resulted in more accurate embeddings for datasets with these characteristics (van der Maaten and Hinton 2008; van der Maaten 2009; van der Maaten and Hinton 2012; van der Maaten 2014). Recently, *Zhu et al.* (2018) were able to predict 3D structures of individual chromosomes using a manifold-learning approach (similar to t-SNE) combined with multi-conformation optimization. Their tool was shown to outperform many of the existing MDS-based methods but could not be applied to the entire 3D-GRP due to the underlying time complexity of multi-conformation optimization (Zhu *et al.* 2018). Based on the results of this regional prediction tool, t-SNE should result in more accurate solutions to the 3D-GRP when compared to existing MDS methods. To test this hypothesis, we developed a new workflow called StoHi-C (pronounced "stoic") that uses t-SNE to predict 3D genome structure from Hi-C data. StoHi-C and MDS were used to predict 3D genome structure for four existing fission yeast datasets (wild-type, G1-arrested, *rad21* deletion and *clr4D* deletion). Overall, StoHi-C was able to more clearly recapitulate well-documented features of fission yeast chromosomal organization (such as the RabI structure) when compared to the MDS method.

## METHODS

StoHi-C is a two step workflow that involves (1) 3D embedding and (2) visualization. A more detailed description of each step is provided in the subsections below. Each step can be run independently or users can invoke an automated shell script ^1^ that runs each step in succession. Complete documentation describing expected inputs, outputs and software requirements can be found on the project homepage ^2^. In the subsequent sections, the StoHi-C workflow is described in general, but also provides details regarding the specific illustrative examples presented in this paper.

### Step 1: 3D Embedding

The 3D coordinates for each genomic bin are calculated using t-SNE (van der Maaten and Hinton 2008; van der Maaten 2009; van der Maaten and Hinton 2012; van der Maaten 2014). A python script ^3^ was developed that accepts a normalized whole-genome contact map as input and outputs the (*x*, *y*, *z*) coordinates for each genomic bin. An example of the required input and expected output can be found on the project homepage. This script uses the TSNE method from the sklearn.manifold library to embed genomic bins in 3D space. The exact parameter values that were used for the fission yeast datasets as well as a brief description their function follow: n_components = 3 (embedding dimensionality), perplexity=5.0 (number of nearest neighbours), early_exaggeration=3.0 (controls the tightness of clusters), n_iter=5000 (maximum number of iterations), method=‘exact’ (do not use the Barnes-Hut approximation), init=‘pca’ (use a principle component analysis to initialize the embedding). These values were selected based on the suggestions provided on t-SNE’s homepage ^4^.

### Step 2: Visualization

Once the (*x*, *y*, *z*) coordinates are generated a multitude of different tools can be used for visualization. Three options are discussed below but any graphing or network visualization tool that accepts 3D coordinates (where *x*, *y*, *z* values are space-delimited with each point on a separate row) could be used.

1. `plot.ly`: A python script `plotly_viz.py` ^5^ was developed that accepts the (*x*, *y*, *z*) coordinates generated in Step 1 and produces a static PNG image and an interactive 3D graph (HTML) using the `plot.ly` library. (Plotly Technologies Inc 2015). The interactive graph can be opened in any web browser. *This option was used to generate the figures for the illustrative examples in this manuscript*.
2. `matplotlib`: A python script `matplotlib_viz.py` ^6^ was developed that accepts the (*x*, *y*, *z*) coordinates generated in Step 1 and produces a static `PNG` image of the corresponding 3D graph as well as a simple `MP4` animation that rotates around the y-axis. This script uses the `py.plot` and animation modules from the `matplotlib` library (Hunter 2007) as well as the `mpl_toolkits.mplot3d` toolkit ^7^.
3. `Chart Studio`: Alternatively, `plot.ly` has a web-based, interactive version available online called Chart Studio ^8^. The 3D coordinates can be directly uploaded to the website to generate an interactive graph. Chart Studio has provided a detailed tutorial on generating this type of visualization ^9^. Customized styles such as colours, labels, size, transparency, etc. for nodes and/or edges can then be set directly by the user through the graphical user interface. Additional node and/or edge attributes can be added to the 3D graph to incorporate complementary biological datasets (if available) with the visualization.

Options 1 and 2 have been automated for the visualization of fission yeast datasets with 10kb resolution. Applying them to datasets from other organisms or datasets with different resolutions would require slight adjustments to the scripts. Documentation of how to make these changes is provided on the project homepage ^10^. Option 3 can not be automated since it is a graphical user interface.

### Comparison with MDS

In order to compare the results of StoHi-C with MDS, the generation of (*x*, *y*, *z*) coordinates in step 1 was also done with metric-MDS. The use of metric-MDS for 3D genome prediction has been widely used since 2010 (*Duan et al.* 2010; *Tanizawa et al.* 2010). Similarly to Step 1 of the StoHi-C workflow, a python script was developed ^11^ that accepts a normalized whole-genome contact map as input and outputs the (*x*, *y*, *z*) coordinates for each genomic bin. This script uses the `MDS` method from the `sklearn.manifold` library (*Pedregosa et al.* 2011) to embed genomic bins in 3D space. The exact parameter values that were used for the fission yeast datasets as well as a brief description of their function follow: `n_components = 3` (embedding dimensionality), `metric=True` (use metric MDS), `max_iter=5000` (maximum number of iterations), `dissimilarity=‘precomputed’` (use a custom dissimilarity matrix). To be consistent with the StoHi-C workflow, the `plot.ly` script used for Step 2 (described above) was used to visualize the results for the illustrative examples in this manuscript.

### Data Availability

The datasets supporting the conclusions of this article were originally generated by *Mizuguchi et al.* (2014) and are available in the Gene Expression Omnibus database (accession number: GSE56849 ^12^). The specific sample numbers are *999a* wild-type (GSM1379427), G1-arrested (GSM1379429), *rad21-K1* mutation (GSM1379430) and *clr4* deletion (GSM1379431).

### Web Resources

StoHi-C is freely available at https://github.com/kimmackay/StoHi-C and is licensed under the Creative Commons Attribution-NonCommercial-ShareAlike 3.0. It requires Python3 and local copies the following libraries: numpy, `sklearn` and `plot.ly`. These libraries are open access and can be downloaded through a package manager like pip or conda. Archived versions of the scripts used to generate the results in this manuscript are available as supplemental data.

## RESULTS & DISCUSSION

The StoHi-C workflow and MDS method described above were used to generate 3D genome predictions for four existing Hi-C fission yeast datasets (*999a* wild type, G1-arrested, *rad21-K1* mutation, and *clr4* deletion) (Mizuguchi *et al.* 2014). Depending on the method, either t-SNE or MDS was used to generate (*x*, *y*, *z*) coordinates. These results were then visualized with `plot.ly` which generates both static images and interactive graphs. Images representing the genomic predictions for each dataset with the StoHi-C workflow (Panels A,C,E,G) and MDS method (Panels B,D,F,H) are presented in Figure 1. Interactive versions of each `plot.ly` graph can be found on the project homepage ^13^. Additionally, the project homepage contains the resultant images and animations generated with the `matplotlib` visualization for the *999a* wild-type dataset ^14 15^.

**Figure 1.**
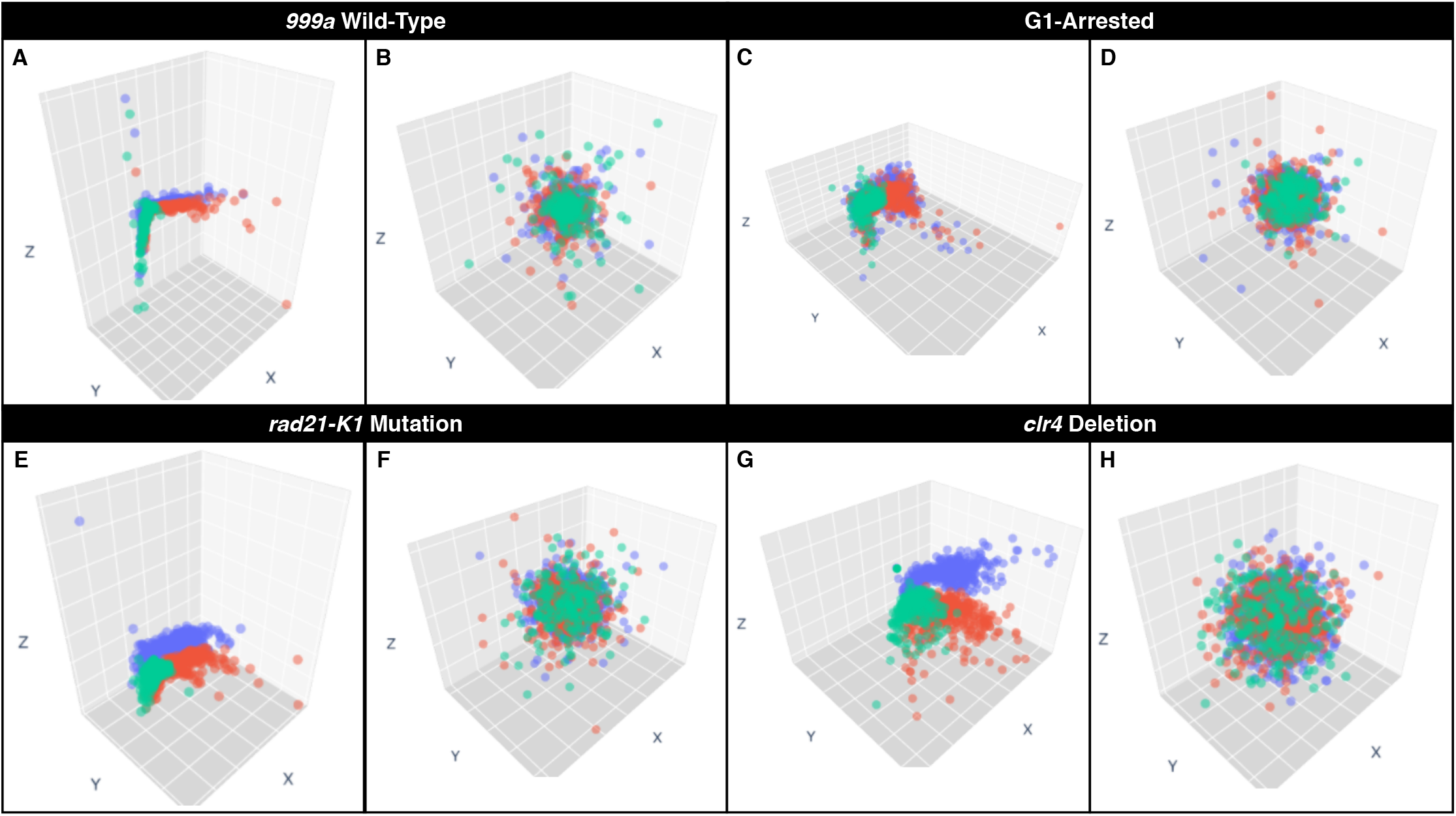
Visualizations of 3D genome predictions for four fission yeast datasets using StoHi-C and MDS. Panels A,C,E and G show 3D genome predictions produced with the StoHi-C workflow, while Panels B,D,F, and H depict the 3D genome predictions generated with MDS. In all panels, chromosomes are indicated with the following colours: purple (chromosome 1), orange (chromosome 2), green (chromosome 3). Corresponding datasets are indicated in the black box directly above the panels (*999a* wild-type: Panels A and B, G1-arrested: Panels C and G, *rad21-K1* mutation: Panels E and F, *clr4* deletion: Panels G and H). In each panel, the *X*, *Y* and *Z* axes are indicated with a corresponding label.

Based on the results presented in Figure 1, the 3D genomic predictions generated with StoHi-C more clearly represent known features of fission yeast chromosomal organization when compared to the MDS method. The StoHi-C predictions (Figure 1A,C,E,G) all clearly depict universal hallmarks of genome organization (e.g. chromosome territories (Cremer and Cremer 2010)) as well as fission yeast specific features (e.g. RabI configuration (Mizuguchi *et al.* 2015; Fernández-Álvarez and Cooper 2017)). Meanwhile, the MDS predictions all resulted in a hairball-like configuration with no apparent biological significance (Figure 1B,D,F,H). This is likely due to a fundamental difference in the algorithms underlying t-SNE and MDS. One of the goals of t-SNE is to preserve the local structure of high-dimensional datasets by placing similar features close together in the final embedding (van der Maaten and Hinton 2008; van der Maaten 2009; van der Maaten and Hinton 2012; van der Maaten 2014). MDS does the opposite, focusing on placing dissimilar features further away in the embedding (Hout *et al.* 2013).

StoHi-C has a worst-case time complexity of *O*(*N*^2^) (the time complexity of t-SNE (van der Maaten and Hinton 2008)) where *N* is the number of genomic bins. This can be improved to *O*(*N*log*N*) by using the Barnes-Hut approximation (van der Maaten 2014) which may be necessary for larger datasets. Classical metric-MDS has a worst-case time complexity of *O*(*N*^3^) (Yang *et al.* 2006) suggesting StoHi-C’s runtime would be better than MDS-based methods in the extreme worst case. Table 1 lists the elapsed runtime required to embed and visualize each dataset using both the StoHiC workflow and MDS method. These timings do not represent a comprehensive complexity analysis and instead are presented to provide context as to whether or not these methods are practical for Hi-C-sized datasets. Interestingly, the MDS embedding is much faster than the t-SNE embedding with average elapsed times of 0.53 seconds and 11.0 seconds, respectively. This could be due to efficiencies in the implementations of the two algorithms.

**Table 1.** Elapsed Runtimes for 3D genome prediction with the StoHi-C workflow and MDS method. Elapsed runtimes are shown in seconds for the embedding (step 1), visualization (step 2) and complete workflow (total).

| Dataset | StoHi-C<br>Elapsed Time |  |  | MDS<br>Elapsed Time |  |  |
| --- | --- | --- | --- | --- | --- | --- |
|  | Step 1 | Step 2 | Total | Step 1 | Step 2 | Total |
| 999a Wild-Type | 10.6 | 5.7 | 16.2 | 0.54 | 5.7 | 6.2 |
| G1-Arrested | 11.7 | 6.3 | 18.0 | 0.52 | 5.9 | 6.4 |
| *rad21-K1* Mutation | 10.8 | 6.3 | 17.1 | 0.54 | 5.8 | 6.3 |
| *clr4* Deletion | 10.7 | 5.8 | 16.5 | 0.51 | 5.6 | 6.1 |

To the best of our knowledge, none of the existing methods for predicting 3D genomic organization have been successfully applied to the datasets used in this paper. Previously, Tanizawa *et al.* (2010) applied MDS to chromosome conformation capture data from fission yeast but the results were not able to recapitulate the RabI configuration of fission yeast chromosomes. StoHi-C was able to produce 3D genomic predictions that are consistent with the large body of work depicting fission yeast genomic organization including the RabI configuration. This is the first time the RabI configuration has been successfully predicted from fission yeast Hi-C data. This is surprising when considering the relative simplicity of the fission yeast genome, but more understandable due to existing tools heavy reliance on MDS. It should be noted that polymer modelling of the same datasets was not successful (Mizuguchi *et al.* 2014).

While StoHi-C appears to be working well with data from the haploid organism fission yeast, additional step(s) may be required to apply it to organisms with higher ploidy (diploid, hexaploid, etc.) if the data is not pre-phased. This is because StoHi-C will have to determine which chromosome copy (or copies) contribute to the detected interactions (the ploidy problem). For now, users should preprocess polyploid Hi-C data with existing phasing tools (see review by Browning and Browning (2011)) prior to using StoHi-C. To solve this problem more permanently, future work will focus on extending StoHi-C to include a step that performs phasing. This is something we are actively working toward in the hopes of applying StoHi-C to polyploid organisms. Once this has been completed, it will be deployed as a new version on the project homepage.

In this manuscript, we present a new workflow called StoHi-C (pronounced "stoic") that uses t-SNE to predict 3D genome structure from Hi-C data. Unlike MDS, t-SNE is well-suited for embedding population-based, sparse, high-dimensional data in 3D space. StoHi-C was used to predict 3D genome structures for four fission yeast Hi-C datasets. The results were compared to the 3D genomic structures predicted from the same datasets using a MDS approach. The 3D genomic predictions generated with StoHi-C more clearly represent known features of fission yeast chromosomal organization when compared to the MDS method. Additionally, this is the first time the RabI 3D genomic organization was successfully predicted from fission yeast Hi-C data. Overall, StoHi-C was able to generate 3D genome structures that more clearly exhibit the established principles of fission yeast 3D genomic organization when compared to the MDS results.

## ENDNOTES

1. https://github.com/kimmackay/StoHi-C/blob/master/stohic.sh

2. https://github.com/kimmackay/StoHi-C/

3. https://github.com/kimmackay/StoHi-C/blob/master/step1/tSNE/run_tSNE.py

4. https://lvdmaaten.github.io/tsne/

5. https://github.com/kimmackay/StoHi-C/blob/master/step2/plotly_viz.py

6. https://github.com/kimmackay/StoHi-C/blob/master/step2/matplotlib_viz.py

7. https://matplotlib.org/mpl_toolkits/mplot3d/index.html#matplotlib-mplot3d-toolkit

8. https://chart-studio.plot.ly/create/#/

9. https://plotly.github.io/make-a-3d-scatter-plot/

10. https://github.com/kimmackay/StoHi-C/issues

11. https://github.com/kimmackay/StoHi-C/blob/master/step1/MDS/run_MDS.py

12. https://www.ncbi.nlm.nih.gov/geo/query/acc.cgi?acc=GSE56849

13. https://github.com/kimmackay/StoHi-C/tree/master/interactive_visualizations/

14. https://github.com/kimmackay/StoHi-C/tree/master/step2/tSNE_results/matplotlib/999a_WT

15. https://github.com/kimmackay/StoHi-C/tree/master/step2/MDS_results/matplotlib/999a_WT

## ACKNOWLEDGEMENTS

This work was supported by the Natural Sciences and Engineering Research Council of Canada [RGPIN 37207 to AK, Vanier Canada Graduate Scholarship to KM].

## SUPPLEMENTAL DATA

The following scripts are archived versions of the scripts used to generate the results presented in this manuscript. For the most recent version and/or to report any software problems please see the project homepage at https://github.com/kimmackay/StoHi-C/

### Supplementary Script 1: StoHi-C

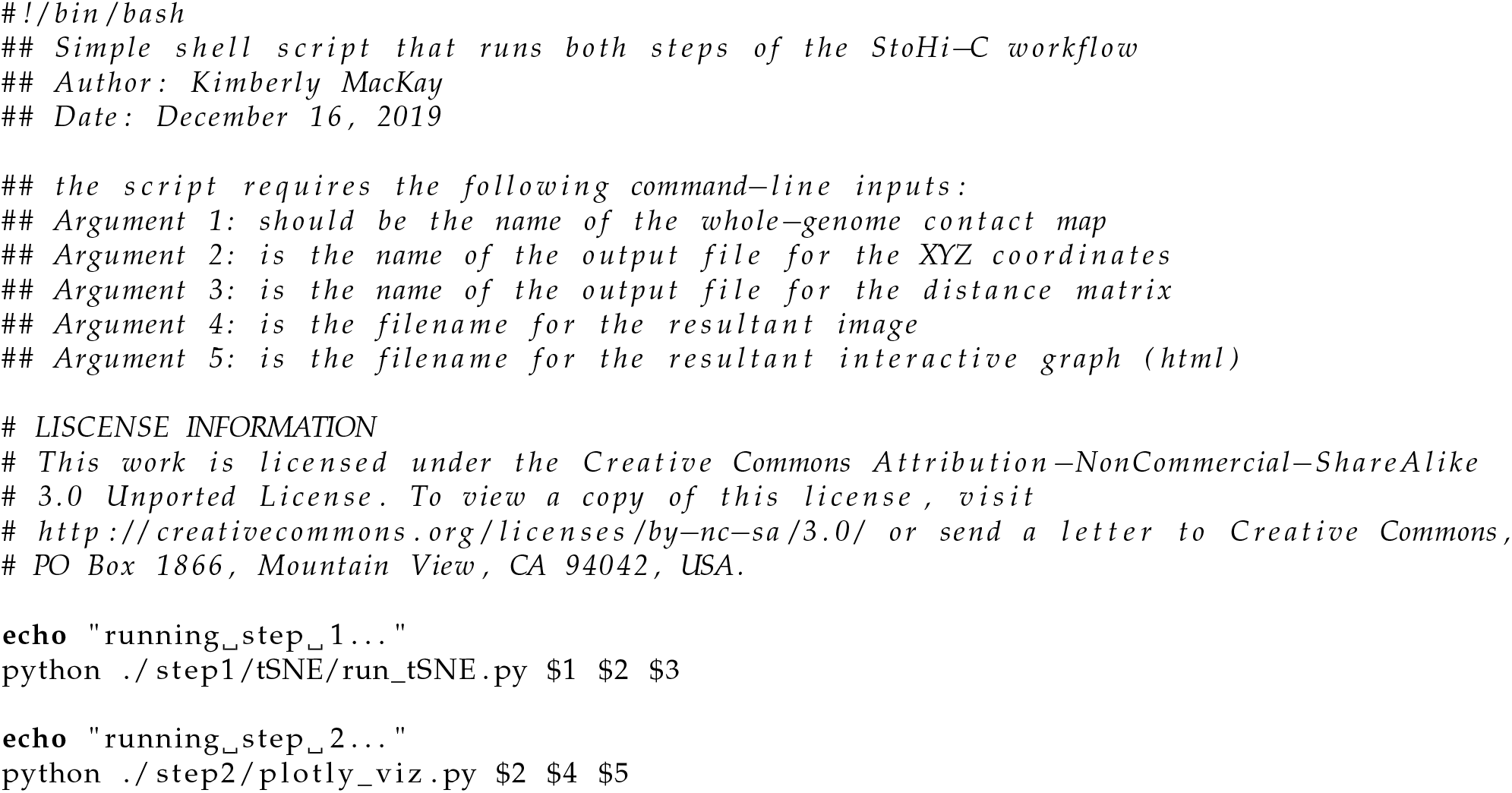

### Supplementary Script 2: StoHi-C Step 1

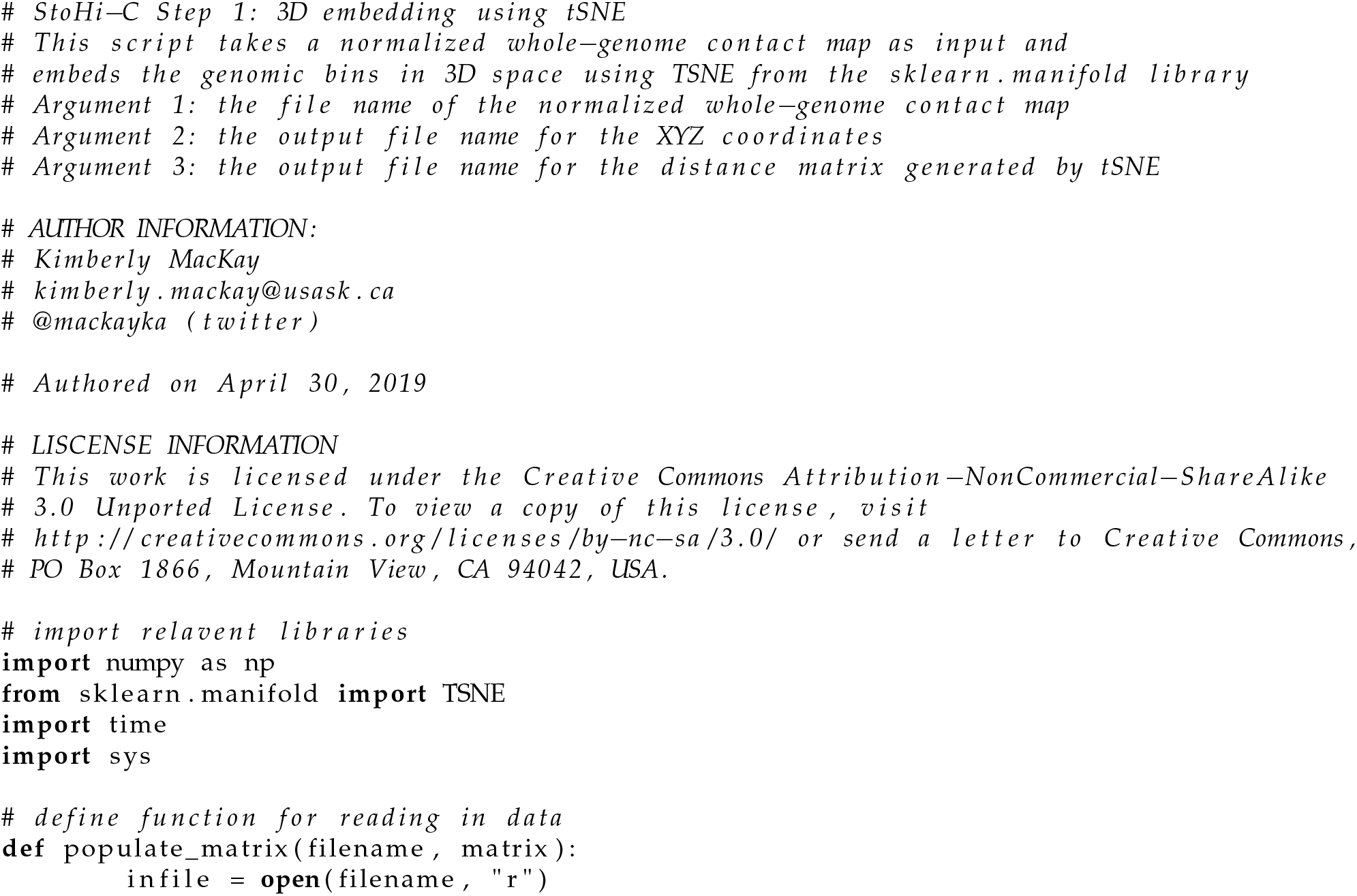
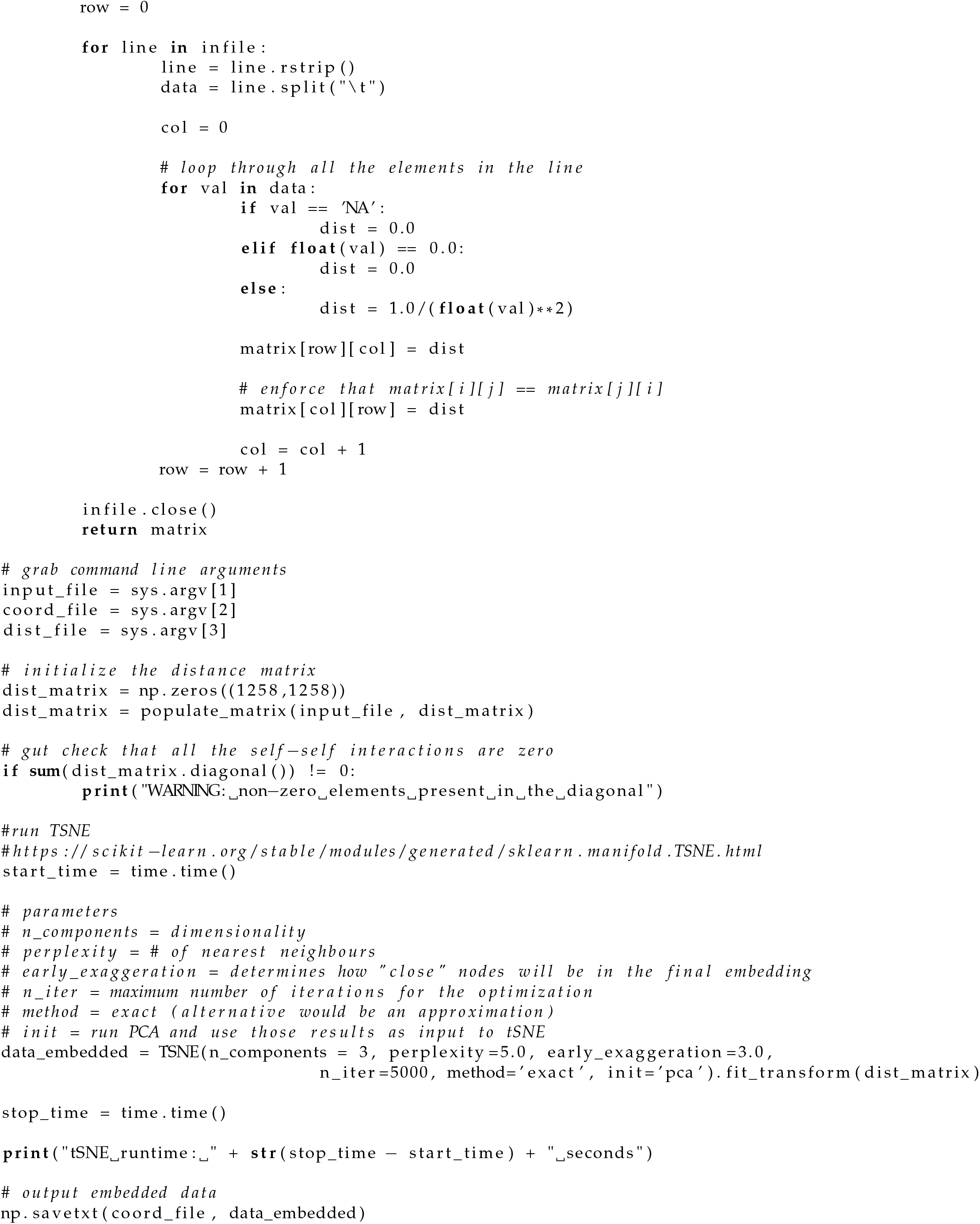
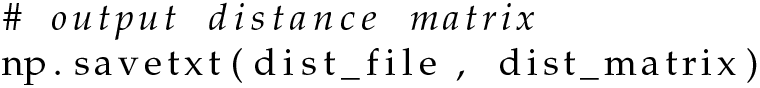

### Supplementary Script 3: StoHi-C Step 2

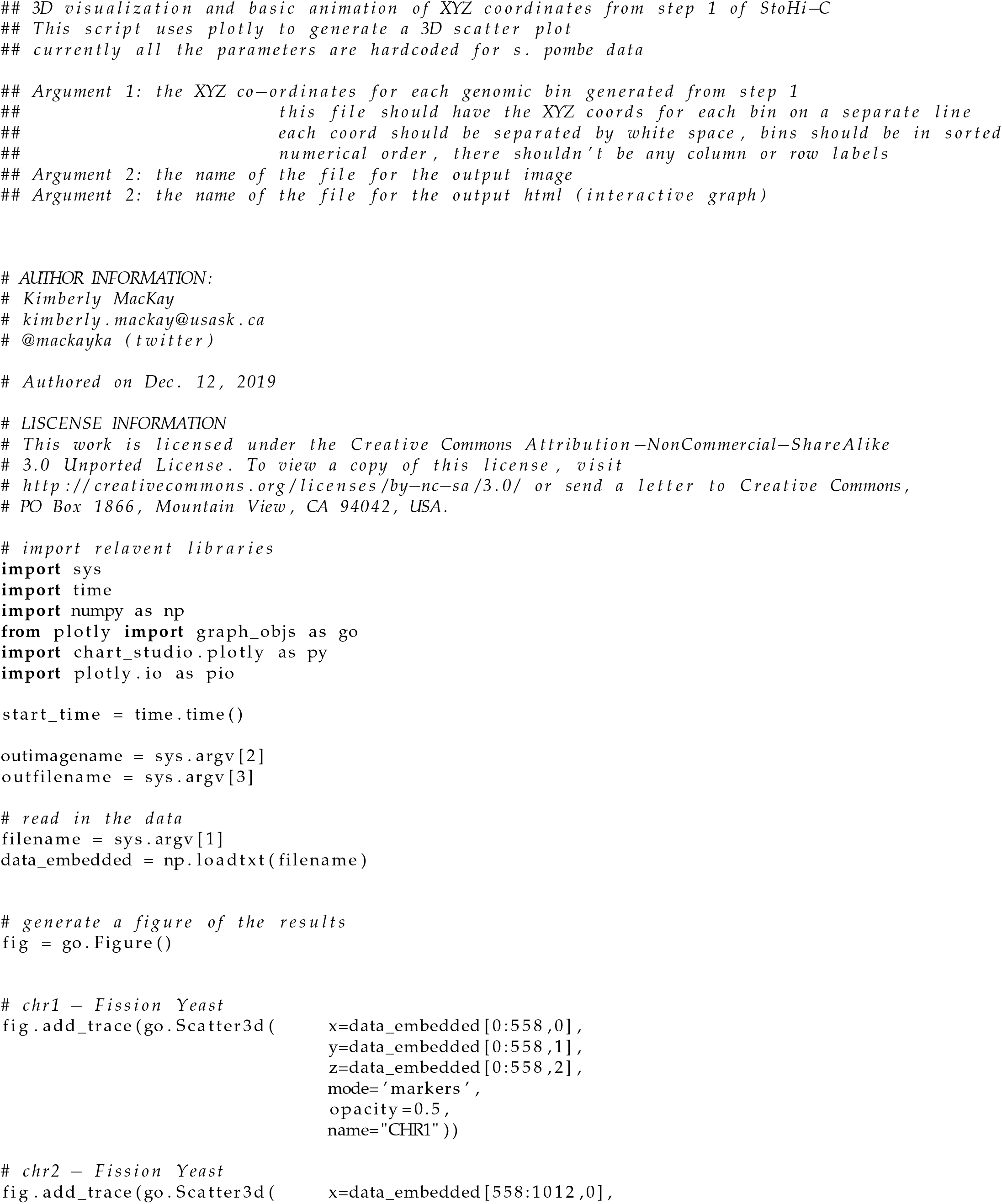
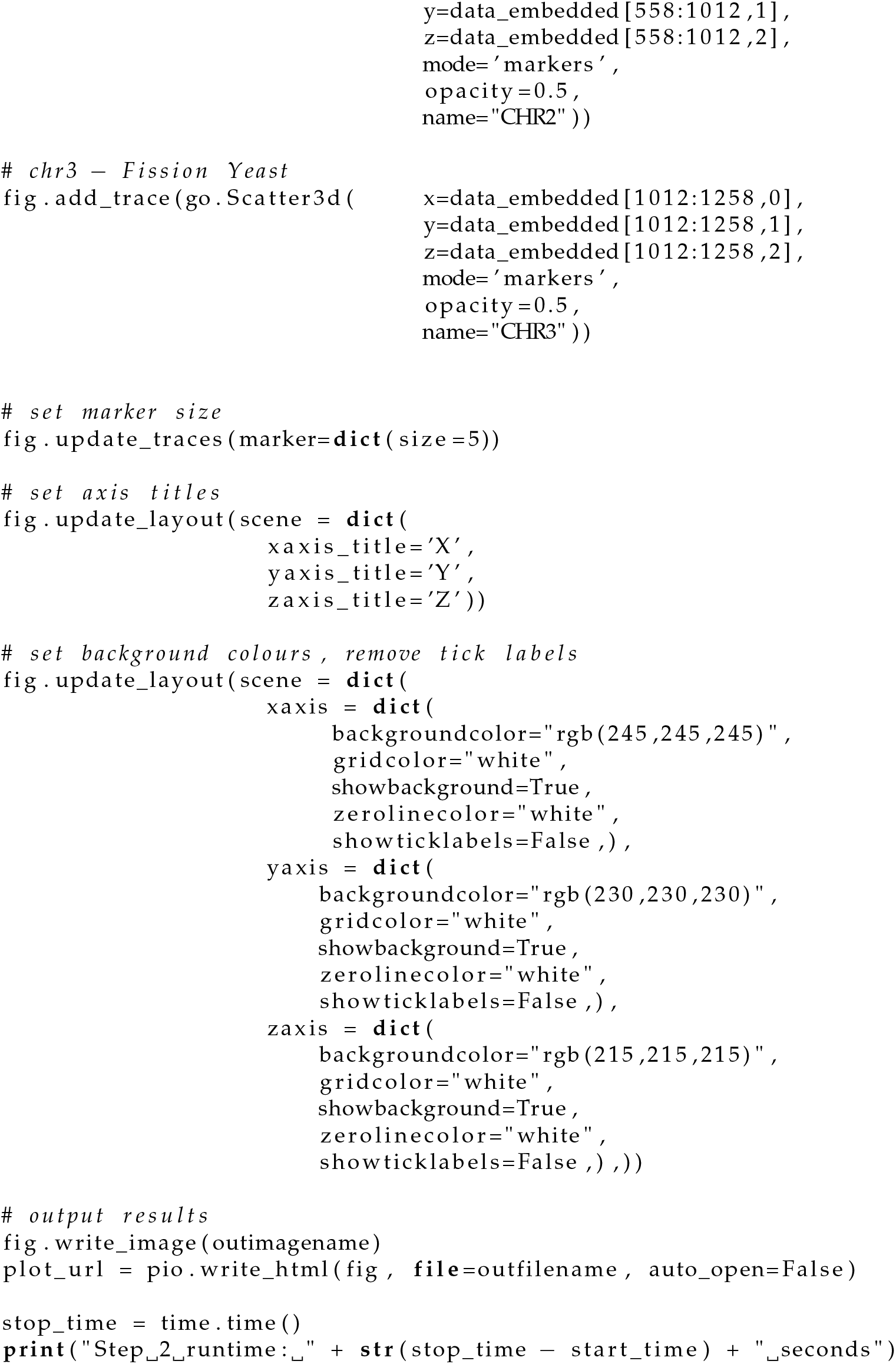

### Supplementary Script 5: MDS

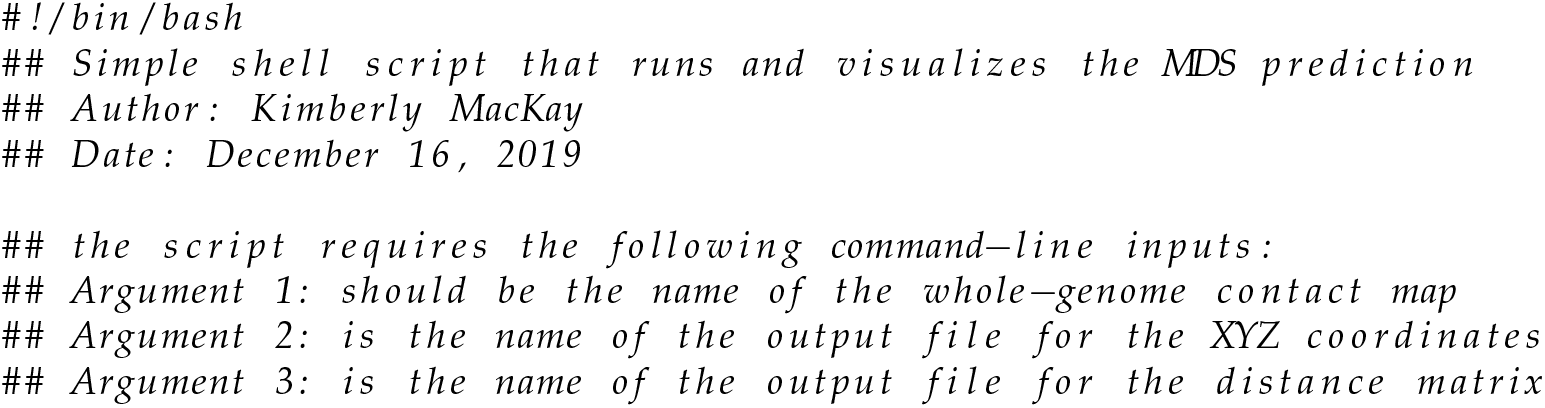
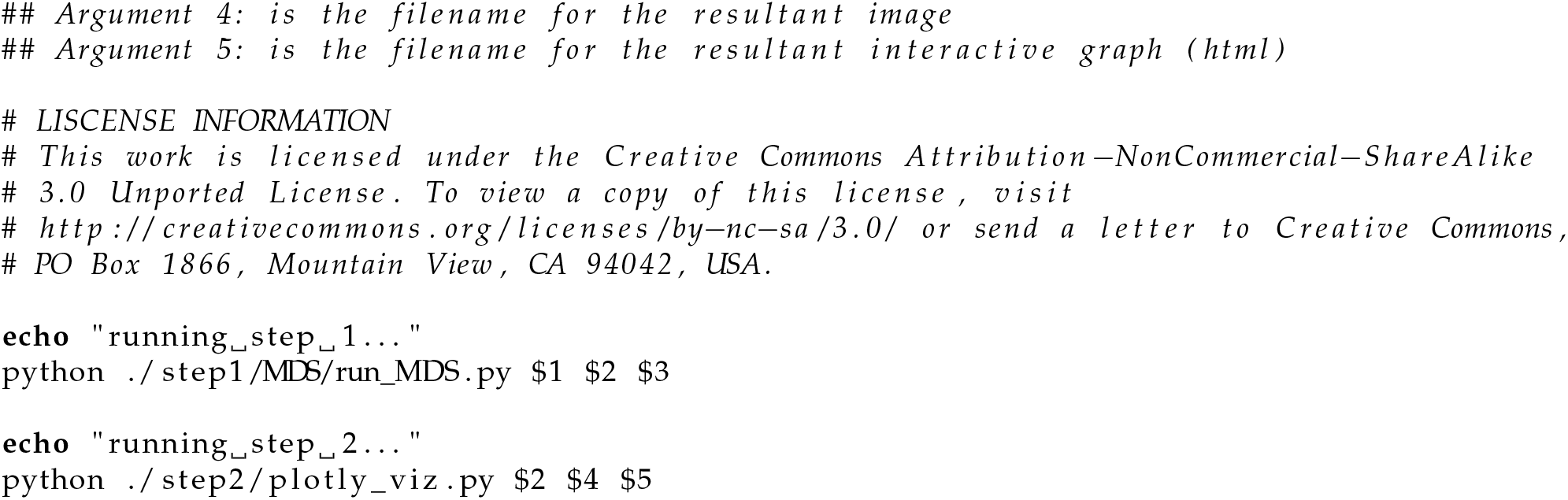

### Supplementary Script 5: MDS Step 1

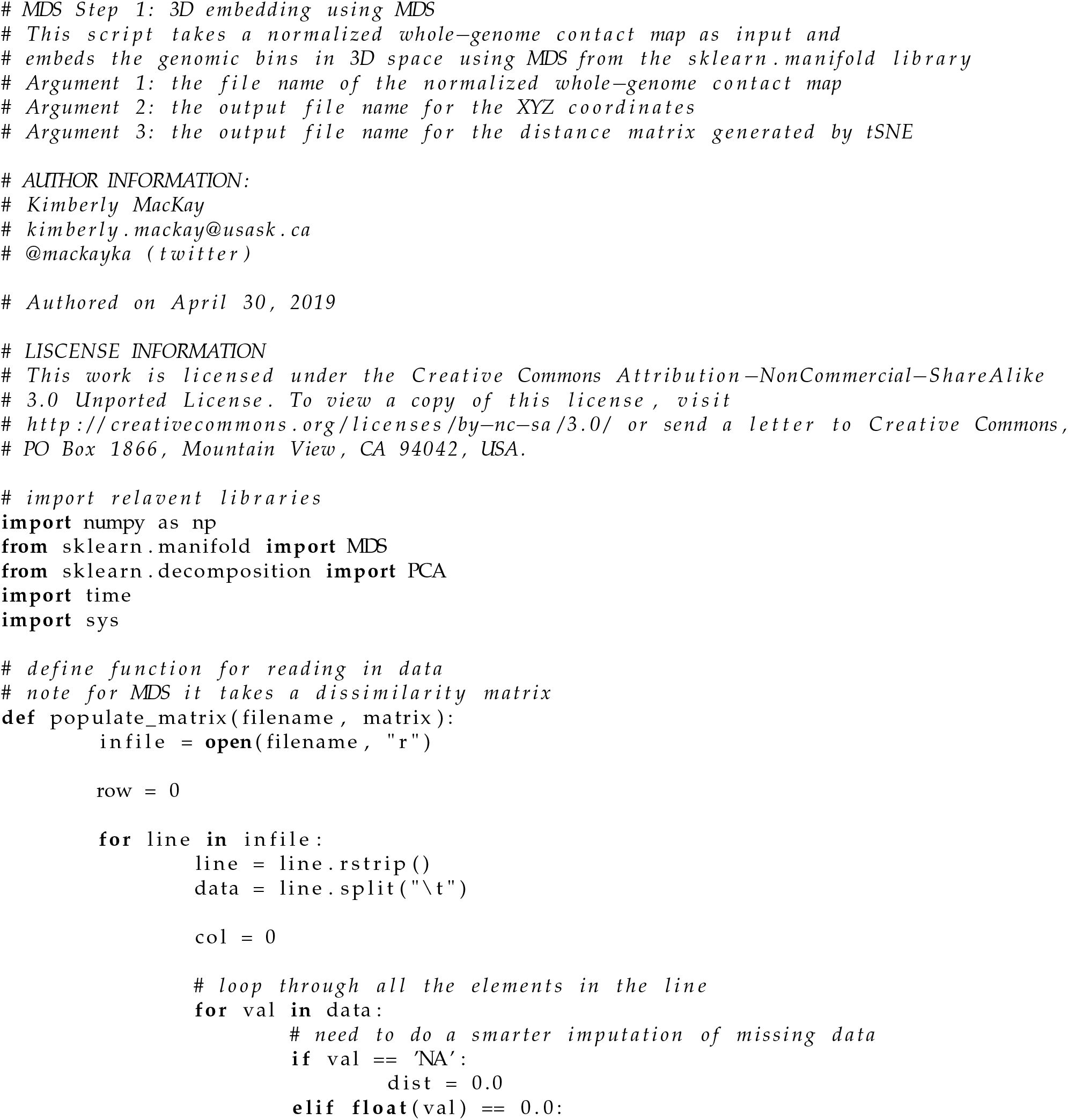
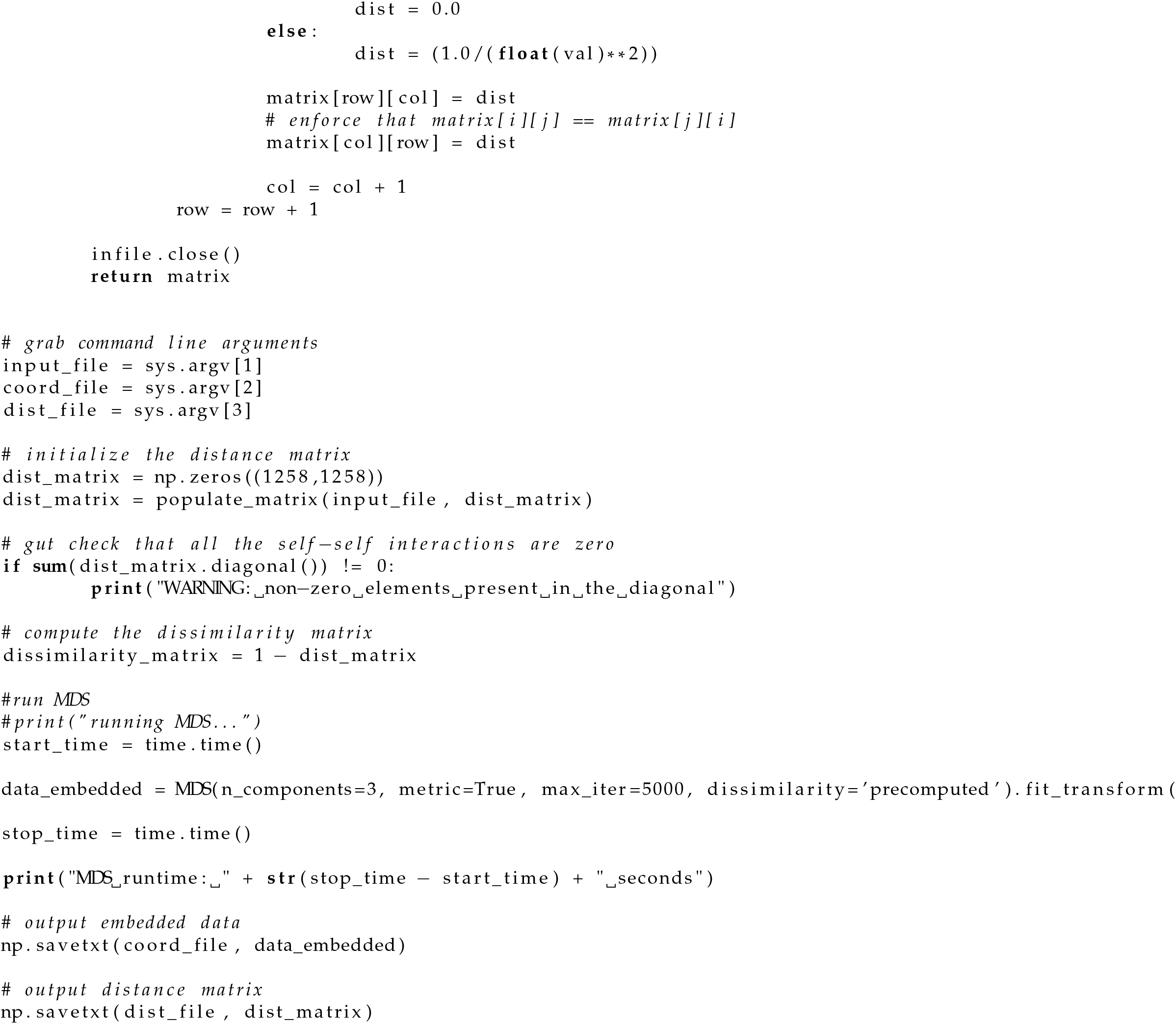

## Notes

https://github.com/kimmackay/StoHi-C/

https://www.ncbi.nlm.nih.gov/geo/query/acc.cgi?acc=GSE56849

